# Enhanced skin permeation of a novel peptide via structural modification, chemical enhancement, and microneedles

**DOI:** 10.1101/2020.10.07.312850

**Authors:** Jungen Chen, Junxing Bian, Basil M. Hantash, David E. Hibbs, Chunyong Wu, Lifeng Kang

## Abstract

Hyperpigmentation is a common skin condition with serious psychosocial consequences. Decapeptide-12, a novel peptide, has been found to be safer than hydroquinone in reducing content of melanin, with efficacy up to more than 50% upon 16 weeks of twice daily treatment. However, the peptide suffers from limited transcutaneous penetration due to its hydrophilicity and large molecular weight. Therefore, decapeptide-12 was modified by adding a palmitate chain in an attempt to overcome this limitation. We also tested the effects of chemical penetration enhancers and microneedles to deliver two peptides through skin. Enhanced skin permeation was found using an *in vitro* human skin permeation model. Moreover, we examined peptide retention of different formulations in skin. Our data showed that palm-peptides in microneedle patch was the most effective.

## Introduction

Melanin, the end product of melanogenesis, plays a crucial role in the absorption of free radicals generated within the cytoplasm, in shielding the host from various types of ionizing radiations, and determining the color of human skin, hair, and eyes [1, 2]. However, excess production of melanin can cause skin hyperpigmentation, a common and non-life-threatening disorder. Melasma is a form of hyperpigmentation that causes brown or gray patches on skin, primarily in the facial area. Hydroquinone and tretinoin, in combination with topical corticosteroids are well established therapeutic agents for melasma and hyperpigmentation treatment [3]. *Hantash* and his group reported that a novel proprietary synthetic peptide, namely, decapeptide-12, which demonstrated a stronger competitive inhibition effect on both mushroom and human tyrosinase enzymes than hydroquinone [1]. Additional studies on humans demonstrated the therapeutic effect of Lumixyl™, a topical formulation containing decapeptide-12, for the treatment of melasma [4, 5].

While decapeptide-12 could reduce production of melanin with good efficacy in treating melasma, the skin permeation of decapeptide-12 remains unclear. As shown in **Fig. 1a**, decapeptide-12 is a polar molecule with multiple amino and hydroxyl groups, which may hinder its transdermal absorption through the lipophilic stratum corneum (SC) [6]. To this end, multiple options are available, e.g., molecular modification [7], using chemical penetration enhancers (CPEs) [8] or microneedles (MNs) [9].

**Figure 1.**
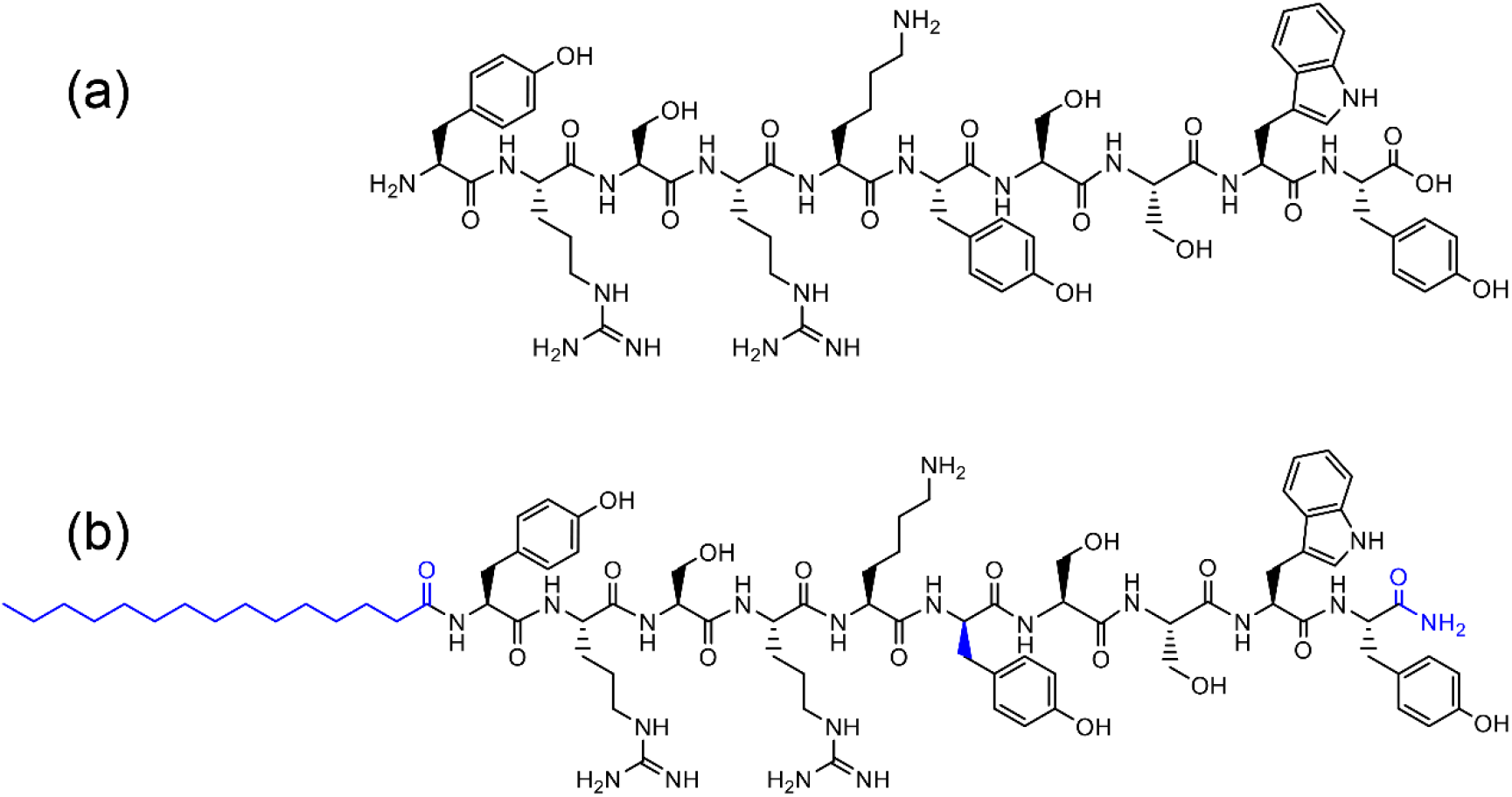
Molecular structures of the two peptides: native peptide (**a**) and its analogue, palm-peptide (**b**). For palm-peptide, N-terminal was palmitoylated, C-terminal was modified to amide, and tyrosine at position 6 was changed from *L*- to *D*-.

For molecular modification, many studies are focused on improving the biostability and bioavailability of peptides for oral, pulmonary, and nasal administration [10]. Due to the lipophilic nature of the SC barrier, peptides existing in zwitterionic form can only permeate minimally through skin. To this end, molecular modification of peptide by increasing its lipophilicity may enhance its skin permeation. In a previous study, we have shown that enhanced skin permeation can be achieved by modifying the structure of a hexapeptide to make it more lipophilic [7].

Chemicals can interact with the intercellular lipids inside SC to enhance drug permeation through skin [11]. Many CPEs, such as fatty acids and terpenes, play an important role in promoting drug permeation through skin [12]. It has been reported that the lipophilicity of the CPEs is proportional to their enhancing effects [8].

In addition to chemical enhancement, there are also physical methods, e.g., electroporation, iontophoresis, diamond microdermabrasion, laser radiation, and ultrasound [13–18]. But disadvantages such as high costs have limited their application. Recently, MNs [19] have emerged as a powerful tool to enhance the skin permeation of a variety of biomolecules, including oligonucleotides, peptides, proteins, and vaccines [20]. MNs can be applied to the skin surface to create an array of microscopic passages through which the biomolecules can reach the dermis [21]. Previously, we reported the application of microneedles on transdermal delivery of lidocaine [22], copper-peptide [23], and nucleic acids [24].

In this study, we investigated the skin permeation of decapeptide-12 (the native peptide, **Fig. 1a**) and its analogue (the palm-peptide, **Fig. 1b**) with higher lipophilicity, on their ability to permeate via skin using both CPEs and MNs. We hypothesize that increased lipophilicity can increase peptide skin permeation, which, however, needs further chemical and/or physical enhancement to get therapeutic effect. Propylene glycol (PG) is a common cosmetic vehicle and shows synergistic action when used together with other CPEs [25]. Oleic acid, camphor, and menthol are widely used CPEs [26–28]. The efficacy of different peptide delivery systems was determined using human cadaveric skin samples.

## Methods

### Materials

Peptide and palm-peptide were synthesized by Bio Basic (Ontario, Canada). PG and PBS tablets were purchased from VWR, USA. Oleic acid, penicillin-streptomycin, and trifluoroacetic acid were purchased from Sigma-Aldrich, USA. Camphor powder was purchased from New Directions, Australia. Menthol crystals was purchased from WFmed, USA. Phosphoric acid solution (85 wt. % in water) was purchased from SAFC, USA. Acetonitrile was obtained from RCI labscan, Thailand. Methanol was obtained from Honeywell, USA. Human epidermis was provided by Science Care, USA. All materials were used as supplied without further purification.

### Peptide synthesis and characterization

The peptide and analogue were synthesized by Bio Basic Canada Inc., using fluorenylmethyloxycarbonyl chloride (FMOC) based synthetic methodology on Rink-amide resin with modifications specific to each compound (**Fig. 2**). The side-chain protecting groups used were: Tyr(tbu), Trp, Ser(tbu), Lys(Boc), Arg(Pbf), Palm. The synthesis was performed using FMOC-Tyr(TBU)-2-Chlorotrityl resin for native-peptide and Rink amide AM resin for palm-peptide, respectively.

**Figure 2.**
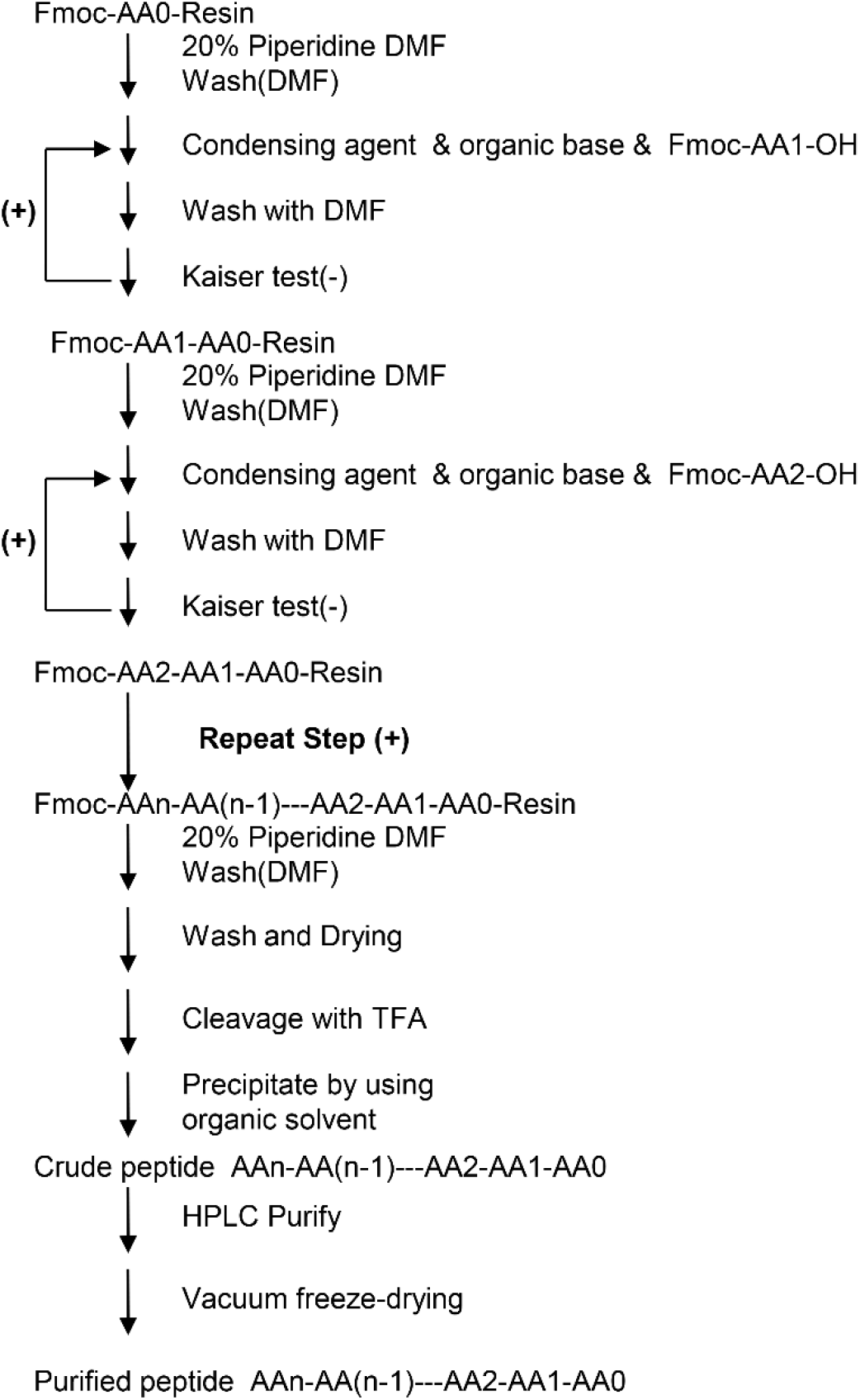
The flow chart of peptide synthesis process.

An initial deprotection of bubbling nitrogen through dimethylformamide (DMF) containing 20% piperidine was followed by a 5-min wash in DMF for 5 times. Conjugation reaction was performed in 500 ml DMF containing 19.3 g 2-(1H-benzotriazole-1-yl)-1,1,3,3-tetramethylaminium tetrafluoroborate (TBTU) and 18.6 ml N,N-diisopropylethylamine (DIEA). The product was washed 3 times using DMF after drying. The whole process was repeated until the conjugation of all the protected amino acids are finished. After conjugation reaction, the on-resin peptide was washed with methanol, dichloromethane, and methyl alcohol sequentially. Drying and lyophilizing process were then carried out. Cleavage was performed using 1000 ml of reagent (82.5% trifluoroacetic acid + 5% 1,2-ethanedithiol + 5% water + 5% p-cresol + 2.5% thioanisole) for 2 h. Afterwards, the resin particles were filtered using a sand-core funnel, for the coarse product to precipitate in the filtered liquid.

The crude peptide was purified using HPLC with a silica gel C-18 column (10 μm). The mobile phase consisted of mobile phase A (Acetonitrile) and mobile B (0.1% trifluoroacetic acid in water) with a gradient elution program with ratios of phase A and B as follows: 76 – 56% B for 0 - 120 min. The flow rate was 200 mL/min. Ultraviolet detection was performed at 220 nm. The sample with a purity higher than 80% was collected, which then was concentrated with the temperature around 35~40 °C. Followed purification, the peptide solution was lyophilized for 24 h in - 40 °C.

### Peptide analogue physicochemical properties

*In silico* prediction of physicochemical properties of two peptides were performed using the Qikprop v6.2 package, which is part of the Schrödinger Suite of software (Schrödinger Release 2020-3: QikProp, Schrödinger, LLC, New York, NY, 2019)

### Saturated peptide solution preparation

The solubility of the peptide was determined in PG to prepare saturated peptide solutions. Excess peptide was added into each solvent inside a 2 mL Eppendorf tube which was kept shaking in an incubation orbital shaker (OM15C, Ratek, Australia) for 48 h. The set up was kept at room temperature. Samples were then centrifuged at 12,000 rpm for 10 min. Subsequently, the supernatant was transferred into an amber bottle for assay.

### Human skin membrane preparation

Human dermatomed skin was obtained from Science Care (Arizona, USA). The skin tissues were excised from the thighs of two male cadavers with age at death of66 and 57. The skin samples used in the study were without identifier and exempted from ethical review. The Integrity of cadaver skin was checked using visual inspection before use to ensure no pores or breaks in the skin surface were present. In addition, any compromise in skin integrity was identified through observation of a rapid and large increase in the amount of permeated peptide through the skin at the first sampling time point.

### Fabrication of micromould

Using a previously reported method [29], the molds were fabricated via thermal curing of Polydimethylsiloxane (PDMS) with embedded 3M Microchannel Skin System as the master template. Briefly, the elastomer and curing agent were mixed and vacuumed at 95 kPa for 10-20 min to remove the entrapped air bubbles. Then, the mixture was poured slowly into the plastic petri dish. Later, it was kept for curing in hot air oven at 70 °C for 2 h. The cured PDMS was gently taken out from the petri dish using a surgical blade. The MN master was gently peeled off from the cured PDMS to get the PDMS mold.

### Fabrication of menthol-based MN patches bearing palm-peptide

The PDMS molds were heated on a hot plate at 65 °C for approximately 10 min. While heating the PDMS molds, palm-peptide was melted in cuvettes at 65 °C using a heating block (Major Science) until all contents melted. Two percent palm-peptide was added into menthol and mixed well. PDMS molds were then removed from the hot plate and the mixed materials poured into the cavity. The molds were placed at room temperature and subsequently de-molded when the material solidified.

### In vitro skin permeation testing

Franz type diffusion cells were used. Human epidermis was mounted between donor and receptor compartments and excessive skin at the sides was trimmed off to minimize lateral diffusion. SC was faced towards the donor compartment and the skin area for permeation was 1.327 cm^2^. The receptor compartment was filled with 5.5 mL of 1 mM PBS solution containing 1% (v/v) antibacterial antimycotic solution. Receptor solution was filtered twice to prevent formation of air bubbles beneath the epidermis. Two hundred and fifty μL of solution was added to the donor compartment and covered with wraps. MN patches bearing peptide were first cut into the same proper size. Patches were then applied to the skin using a finger-thumb grip for 20 s. Scotch tape was used to fix the patches onto the skin. Triplicates were prepared. The donor compartment and the sampling port were covered with wraps to minimize gel contamination and evaporation. Magnetic stirrers were used at 180 rpm to mix the receptor solution during the study. The device was kept at 32 °C. Five hundred μL of the receptor fluid were collected at different time points for HPLC analysis and replaced with 500 μL of fresh receptor solution. Permeation was measured over 24 hr.

### HPLC for the peptide analysis

The amount of peptide permeated was determined using Shimadzu CBM-20A HPLC system with Agilent ODS C18 column (4.6 mm x 250 mm x 5 μm, 170Å). The mobile phase consisted of mobile phase A (0.1% v/v trifluoroacetic acid in water) and mobile B (0.1% v/v trifluoroacetic acid in acetonitrile) with a gradient elution program with ratios of solvents A and B as follows: native peptide (15 – 35% B for 0 - 20 min and 15% for 20.01-30 min), palm-peptide (55 – 45% B for 0 - 10 min and 55% for 10.01-30 min) The flow rate was 1.0 mL/min. The injection volume was 50 μL and ultraviolet detection was performed at 280 nm. A calibration curve was established using the standard solutions from 5 μg/mL to 50 μg/mL for native peptide and from 1 μg/mL to 50 μg/mL for palm-peptide.

### Statistical analysis

All data were collated and prepared using GraphPad Prism 8 (GraphPad Software Inc, CA, USA). Statistical significances were calculated using ANOVA, with p < 0.05 considered statistically significant.

## Results

### Peptide modification and physicochemical properties

The structures of the peptides were verified the HPLC and MS data (**Fig.3**). Although the lipophilicity of palm-peptide was increased after modifications, its retention time was shorter than the native peptide. The reason may be because of the ratio of organic phase in the gradient elution program was different, i.e., 20% for the native peptide while 40% for the palm-peptide.

**Figure 3.**
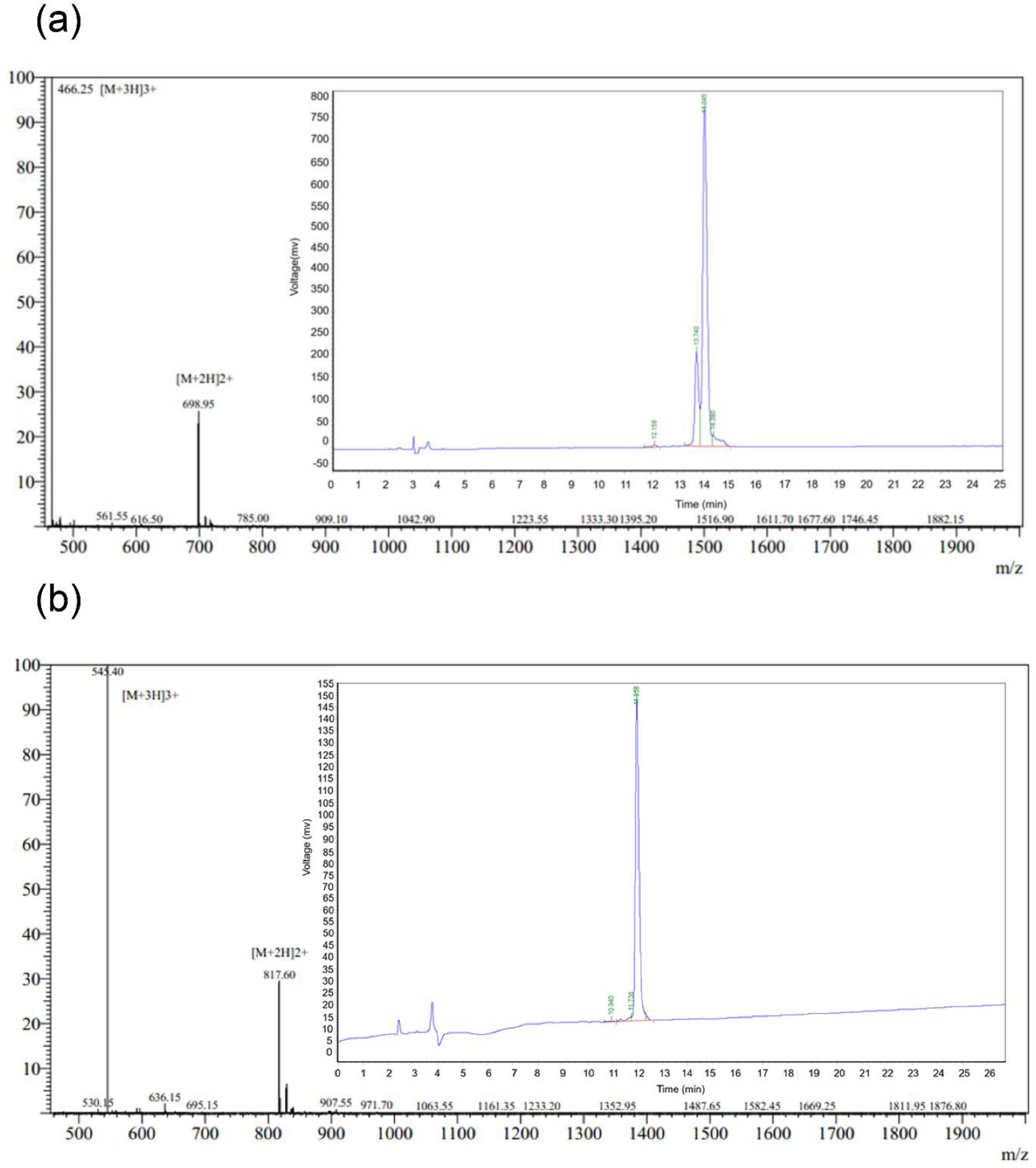
HPLC and MS data for native peptide (**a**) and palm-peptide (**b**).

The higher amount of organic solvent in the mobile phase shortened the retention time of modified peptide, despite the increased lipophilicity. The molecular structure of the peptides was verified by the MS spectra of the [M+3H]+ in m/z 466.25 and [M+2H]+ in m/z 698.95 for native-peptide and the [M+3H]+ in m/z 545.40 and [M+2H]+ in m/z 817.60 for palm-peptide, respectively.

The palm-peptide was modified from its parent compound, namely, the native peptide, to reduce the formation of zwitterions and increase lipophilicity. For palm-peptide, N-terminal was palmitoylated, C-terminal was modified to amide, and the tyrosine at position 6 was changed from *L-* to *D*-. After C-terminal esterification, as seen in palm-peptide, the originally ionizable carboxyl ground no longer formed charged ions. The palm-peptide showed higher LogP of −0.8 than the native peptide, which has a LogP of −6.5 (**Tab.1**). However, both peptides have high molecular weight, making it difficult for them to permeate through skin.

**Table 1.** Physicochemical properties of the peptides by silico prediction.

|  | **Native peptide** | **Palm-peptide** |
| --- | --- | --- |
| *Sequence* | YRSRKYSSWY | YRSRK[*Y]SSWY |
| *Formula* | C<sub>65</sub>H<sub>90</sub>N<sub>18</sub>O<sub>17</sub> | C<sub>81</sub>H<sub>121</sub>N<sub>19</sub>O<sub>17</sub> |
| *LogP* | -6.5 | -0.8 |
| *MW (Da)* | 1395.5 | 1633.0 |
| *H-donors* | 19 | 18 |
| *H-acceptors* | 28 | 29 |
| *Solubility (mg/ml)* | 1000 | 2.6 |

### In vitro skin permeation of peptides

PG is commonly used in cosmetics, well known for its co-solvent and permeation enhancing effect. Hence, pure PG and PG containing 5% (w/v) enhancer were also used to prepare peptide solutions. The amount of native peptide in MN patch and palm-peptide in different drug delivery systems is shown in **Fig. 4a**. The highest permeation of palm-peptide was observed in PG containing 5% (w/v) menthol. After normalization against dose, as shown in **Fig. 4b**, a significant higher percentage of native peptide permeated through skin in the MN patch than in other formulations (P<0.0001).

**Figure 4.**
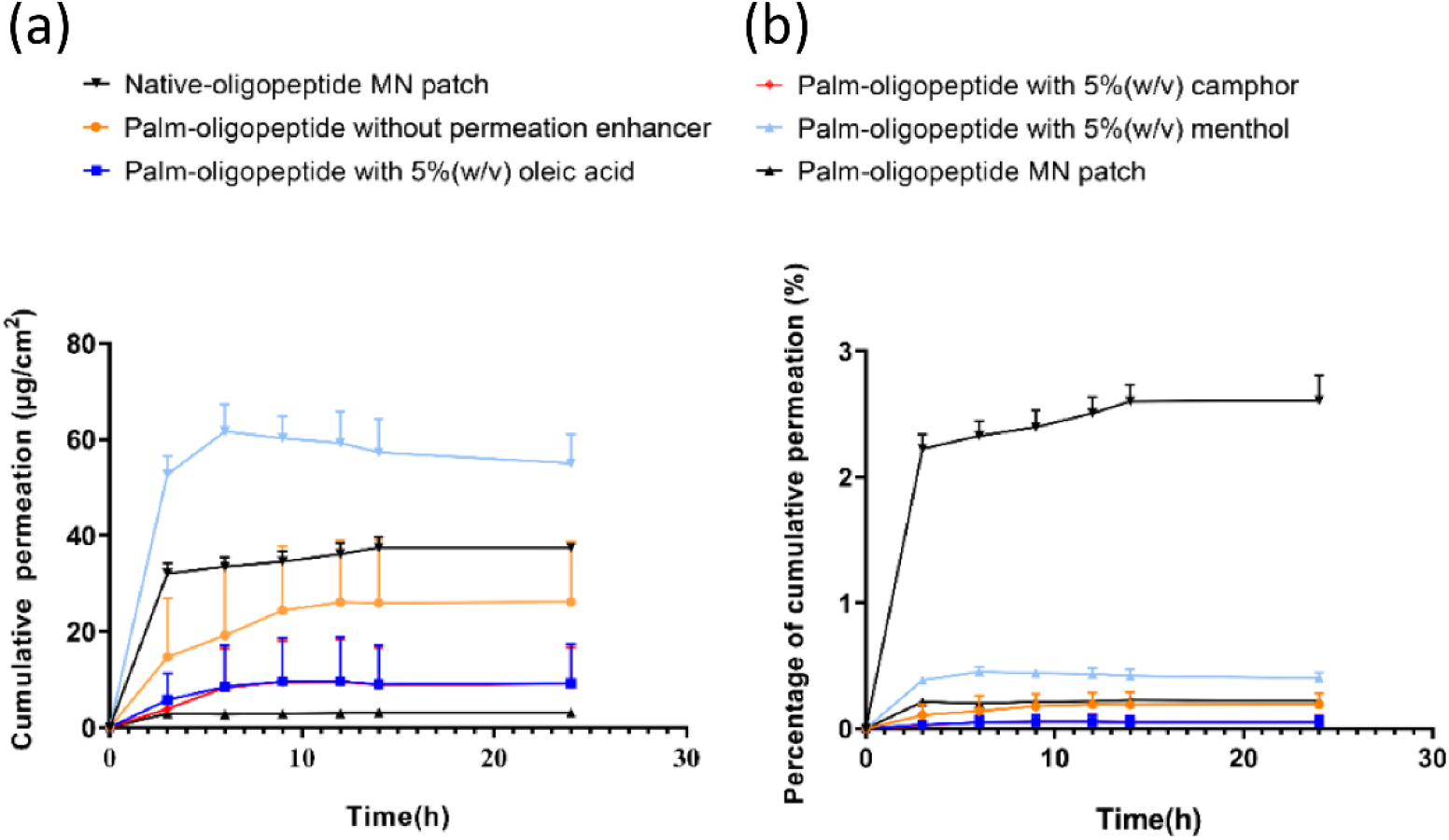
Cumulative permeation profiles of the 2 peptides over 24 h in different carriers. Native peptide in MN patch and palm-peptide in PG, PG with 5% (w/v) oleic acid, PG with 5% (w/v) camphor, PG with 5% (w/v) menthol, and MN patch: amounts (**a**) and percentage (**b**).

### Cumulative peptide permeation in 24 h

The cumulative permeation of the two peptides in 24 h from different carriers is shown in **Fig. 5**. The highest permeation of palm-peptide was found in 5% (w/v) menthol (**Fig. 5a**). For MN patches, a significantly higher amount of native peptide permeated through skin than the palm-peptide. After normalization by dose (**Fig. 5b**), the native peptide permeation in MN patch was significantly higher than others.

**Figure 5.**
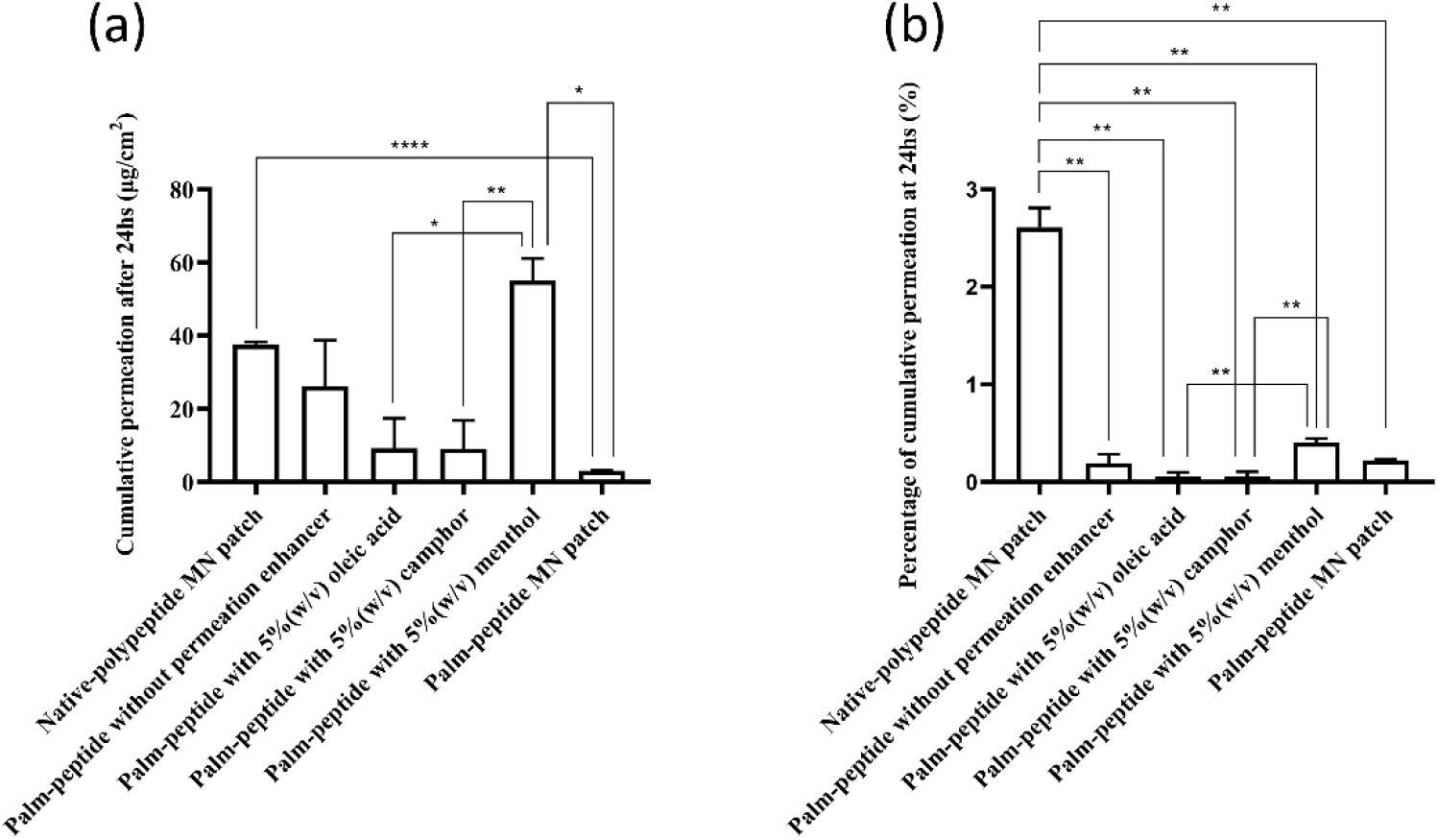
Cumulative amount of peptide permeated through skin after 24 h, in amounts (**a**) and percentage (**b**). (*, P<0.05; **, P<0.01; ***, P<0.001; ****, P<0.0001).

### Peptide human skin retention

Since the peptides reduce epidermal melanin content, we measured skin retention of the two peptides after 24 h (**Fig. 6**). No peptides were detected in the skin samples for the following 3 formulations: native peptide in MN patch, palm-peptide in 5% camphor, and palm-peptide in 5 % menthol. For the rest of the formulations, no significant differences were found among them (**Fig. 6a**). After dose normalization (**Fig. 6b**), the percentage of palm-peptide retained in skin, when using MN patch, was significantly higher than the other two formulations.

**Figure 6.**
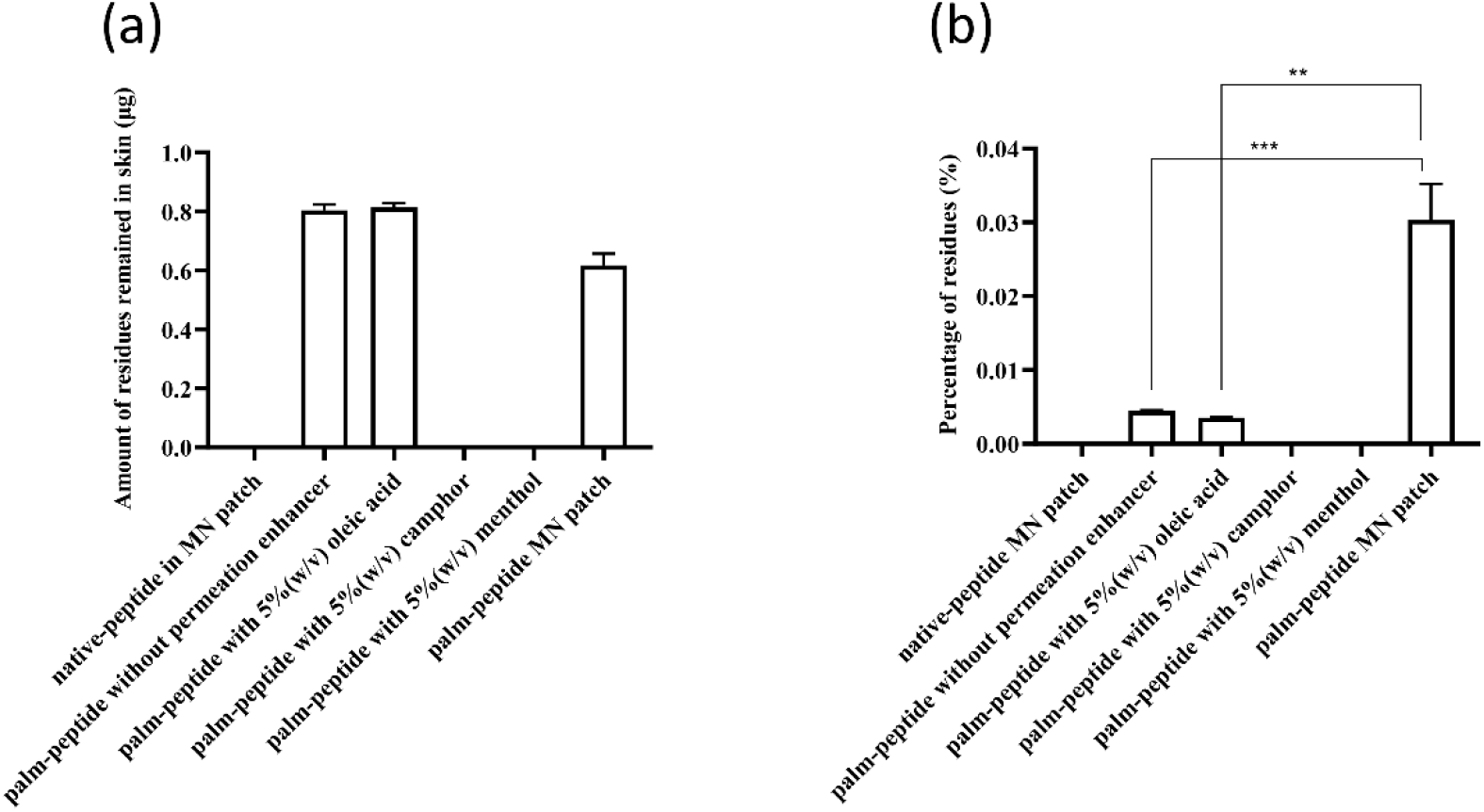
The skin retention of two peptides after 24 h, in amounts (**a**) and percentage (**b**). Retention of native peptide in MN patch, palm-peptide in 5% (w/v) camphor / menthol were not detected. (**, P<0.01; ***, P<0.001).

## Discussion

### The skin barrier and modification of the peptides

The outermost lipophilic layer of the skin, namely, SC, is the major obstacle to percutaneous penetration. This lipophilic layer ensures that only small and moderately lipophilic molecules can traverse the skin in sufficient quantities to elicit therapeutically relevant effects [30]. Peptides with large molecular weight and hydrophilic properties cannot penetrate SC in sufficient amounts. Therefore, we modified the termini of native peptide and synthesized a new peptide, namely, palm-peptide. Because of the higher lipophilicity of palm-peptide than native peptide, there was a corresponding increase in amount of permeation and skin retention in 24 h (**Fig. 5** and **Fig. 6**) in PG solutions. It suggested the feasibility of chemical modification in promoting transdermal delivery of peptides using PG solution.

### The skin retention of the peptides

Since the two peptides target melanocytes to exert their therapeutic effect [1], and melanocytes are located in the basal layer of the epidermis, we assessed the retention of the peptides in side skin membranes. In comparison to undetectable skin retention of native peptide when delivered using MN patches, there was apparent skin retention of palm-peptide when delivered in MN patches (**Fig. 6**). Therefore, structural modification of native peptide can largely increase its efficacy topically by increasing its skin retention.

### The effect of CPEs on skin permeation of the peptides

CPEs have been investigated to promote drug transdermal delivery for decades and many have improved transdermal delivery [31, 32]. One mechanism of action has been described as ‘pull-push’ effect [33]. High solubility parameter difference between CPE and the drug would push the drug molecules through SC. According to the theory and considering the property differences between the two peptides, we selected oleic acid with high lipophilicity and hydrophilic camphor and menthol (**Tab. 2**), which have previously been reported to enhance transdermal penetration [26–28]. We expected PG containing 5% (w/v) oleic acid would exhibit the best improvement of skin permeation and skin retention due to the greatest LogP difference from both peptides.

**Table 2.** The properties of three chemical penetration enhancers.

|  | **Oleic acid** | **Camphor** | **Menthol** |
| --- | --- | --- | --- |
| *LogP* | 7.4 ± 0.2 | 2.1 ± 0.3 | 3.2 ± 0.2 |
| *MW (Da)* | 282.5 | 152.2 | 156.3 |
| *Solubility (mg/ml)* | 0.002 | 1.1 | 1.5 |

However, results were not consistent with the expectation that oleic acid will have the best enhancing effect on skin permeation. For the native peptide, the 3 CPEs didn’t alter skin permeation. For palm-peptide, camphor and oleic acid showed inhibitory effects, while menthol had an enhancing effect on skin permeation. This may be due to the interaction between CPEs and skin leading to altered partition behavior of palm-peptide.

### The effect of MNs on skin permeation of the peptides

In this study, both peptides in MN patches showed more benefits than CPEs, in terms of skin permeation. Moreover, the native peptide showed better skin permeation than palm-peptide. It can be attributed to the higher solubility of native peptide in water than the palm-peptide. MNs destroyed the skin barrier by creating microscale passages through skin, so that native peptides in MN patches can directly transport into the receptor chamber. Native peptide, as a macromolecule with high polarity, tend to diffuse into the receptor PBS solution, in which native peptide is miscible.

### The effect of MNs on skin retention of peptides

For skin retention, the highest percentage of palm-peptide retained inside skin was observed in MN patch, significantly different from other groups, although the percentage of palm-peptide in MN patch retention inside skin was still very low (<0.1%). This may be because MNs destroyed the skin barrier, peptides directly transported into dermis and epidermis. Besides, higher lipophilicity of palm-peptide facilitated transcutaneous penetration.

On the other hand, no native peptides can be detected in the skin samples after 24 h. This may be because the following two reasons. First, the native peptide is more polar than the palm-peptide, so it tends to diffuse into the aqueous solution in the receptor compartment. Second, the amount of peptide inside the MN patch is very limited. As a result, no native peptide was retained in skin, while high amount of native peptide can permeate through skin.

### Estimation of palm-peptide concentration inside skin

The highest of palm-peptide concentration in skin reached 9.0 μg/g, which was close to effective concentration of cell experiments (about 13.95 μg/g, estimated from 10 μM); what’s more, according to cytotoxic test, only when concentration exceeds 100 μM peptides inhibit melanocytes viability and proliferation [1], far exceeding the concentration that can be achieved in the skin. However, there was no evidence for the efficacy of palm-peptides concentration in skin and therefore, more detailed pharmacodynamic experiments need to be performed.

### Infinite dose vs finite dose in the donor compartment

Depletion of permeants in the donor side over the course of the experiment usually results in a reduction in the rate of permeation and an eventual plateau in the cumulative permeation profile. On the other hand, if the permeant is applied as an infinite dose, there is sufficient permeants in the donor side to make any changes in donor concentration negligible [34]. Infinite dose experiments are those with typical dose of more than 10 mg/cm^2^ [35] in the donor compartment.

In this study, MN patch was applied as finite dose condition and exhibited a decreasing rate and a plateau in cumulative permeation profile (**Fig.4**). However, the theory was not applicable to the permeation profile of palm-peptide solutions, which was applied as infinite dosage condition (>13 mg/cm^2^) yet resulted in a rate reduction as well.

### Ideal LogP for transdermal delivery and peptide modification

Previous studies have shown that the ideal LogP for drug penetration is ~ 2 [36]. This represents a large difference between the LogP of palm-peptide (−0.751) and the ideal value. In this study, we modified the native peptide to increase its lipophilicity. Previously, we reported the improvement of peptides penetration across skin by changing functional groups of side chains, which increased lipophilicity from −6.37 of LogP to 1.75 [7]. That may represent a feasible solution.

## Conclusion

In study, we used chemical modification, CPEs and MN patch to improve the skin permeation and retention of peptides. CPEs exhibited a positive effect on skin permeation but did not promote skin retention of palm-peptide. MN patches improved both skin permeation and retention of palm-peptide. For the peptides to target melanocytes located in the epidermal basal layer, structural modification and MN patches could improve their efficacy.

## Acknowledgement

The authors acknowledge the support from the University of Sydney. The research also received funding support from the Escape Therapeutics Inc., USA and the National Natural Science Foundation of China (No. 82074128).

## References

[1] A. Abu Ubeid, L. Zhao, Y. Wang, B.M. Hantash, Short-sequence oligopeptides with inhibitory activity against mushroom and human tyrosinase, J Invest Dermatol, 129 (2009) 2242–2249.

[2] S. Parvez, M. Kang, H.S. Chung, C. Cho, M.C. Hong, M.K. Shin, H. Bae, Survey and mechanism of skin depigmenting and lightening agents, Phytother Res, 20 (2006) 921–934.

[3] H.M. Torok, A comprehensive review of the long-term and short-term treatment of melasma with a triple combination cream, Am J Clin Dermatol, 7 (2006) 223–230.

[4] B.M. Hantash, F. Jimenez, A split-face, double-blind, randomized and placebo-controlled pilot evaluation of a novel oligopeptide for the treatment of recalcitrant melasma, J Drugs Dermatol, 8 (2009) 732–735.

[5] B.M. Hantash, F. Jimenez, Treatment of mild to moderate facial melasma with the Lumixyl topical brightening system, J Drugs Dermatol, 11 (2012) 660–662.

[6] R. Kumar, A. Philip, Modified transdermal technologies: Breaking the barriers of drug permeation via the skin, Trop J Pharm Res, 6 (2007) 633–644.

[7] S.H. Lim, Y. Sun, T. Thiruvallur Madanagopal, V. Rosa, L. Kang, Enhanced skin permeation of anti-wrinkle peptides via molecular modification, Sci Rep, 8 (2018) 1596.

[8] L. Kang, C.W. Yap, P.F. Lim, Y.Z. Chen, P.C. Ho, Y.W. Chan, G.P. Wong, S.Y. Chan, Formulation development of transdermal dosage forms: quantitative structure-activity relationship model for predicting activities of terpenes that enhance drug penetration through human skin, J Control Release, 120 (2007) 211–219.

[9] S.H. Lim, W.J. Tiew, J. Zhang, P.C. Ho, N.N. Kachouie, L. Kang, Geometrical optimisation of a personalised microneedle eye patch for transdermal delivery of anti-wrinkle small peptide, Biofabrication, 12 (2020) 035003.

[10] Z. Antosova, M. Mackova, V. Kral, T. Macek, Therapeutic application of peptides and proteins: parenteral forever?, Trends Biotechnol, 27 (2009) 628–635.

[11] L. Kang, P.C. Ho, S.Y. Chan, Interactions between a skin penetration enhancer and the main components of human stratum corneum lipids - Isothermal titration calorimetry study, J Therm Anal Calorim, 83 (2006) 27–30.

[12] A. Ahad, M. Aqil, K. Kohli, H. Chaudhary, Y. Sultana, M. Mujeeb, S. Talegaonkar, Chemical penetration enhancers: a patent review, Expert Opin Ther Pat, 19 (2009) 969–988.

[13] C.T. Costello, A.H. Jeske, Iontophoresis: applications in transdermal medication delivery, Phys Ther, 75 (1995) 554–563.

[14] C. Gómez, M. Benito, J.M. Teijón, M.D. Blanco, Novel methods and devices to enhance transdermal drug delivery: the importance of laser radiation in transdermal drug delivery, Ther Deliv, 3 (2012) 373–388.

[15] C. Lombry, N. Dujardin, V. Préat, Transdermal delivery of macromolecules using skin electroporation, Pharm Res, 17 (2000) 32–37.

[16] S. Mitragotri, D. Blankschtein, R. Langer, Ultrasound-mediated transdermal protein delivery, Science, 269 (1995) 850–853.

[17] A. Bhatia, J. Hsu, B.M. Hantash, Combined topical delivery and dermalinfusion of decapeptide-12 accelerates resolution of post-inflammatory hyperpigmentation in skin of color, J Drugs Dermatol, 13 (2014) 84.

[18] B.M. Hantash, Enhanced topical uptake of ascorbic acid in fractional photothermolysis-treated ex vivo human skin, J Dermatol Surg Res Ther, 2018 (2018) 20–27.

[19] G. Ma, C. Wu, Microneedle, bio-microneedle and bio-inspired microneedle: A review, J Control Release, 251 (2017) 11–23.

[20] M.R. Prausnitz, Microneedles for transdermal drug delivery, Adv Drug Deliv Rev, 56 (2004) 581–587.

[21] Y.B. Schuetz, A. Naik, R.H. Guy, Y.N. Kalia, Emerging strategies for the transdermal delivery of peptide and protein drugs, Expert Opin Drug Deliv, 2 (2005) 533–548.

[22] H. Kathuria, H. Li, J. Pan, S.H. Lim, J.S. Kochhar, C. Wu, L. Kang, Large size microneedle patch to deliver lidocaine through skin, Pharm Res, 33 (2016) 2653–2667.

[23] H. Li, Y.S.J. Low, H.P. Chong, M.T. Zin, C.-Y Lee, B. Li, M. Leolukman, L. Kang, Microneedle-mediated delivery of copper peptide through skin, Pharm Res, 32 (2015) 2678–2689.

[24] K.J. Koh, Y. Liu, S.H. Lim, X.J. Loh, L. Kang, C.Y. Lim, K.K.L. Phua, Formulation, characterization and evaluation of mRNA-loaded dissolvable polymeric microneedles (RNApatch), Sci Rep, 8 (2018) 11842.

[25] A.C. Williams, B.W. Barry, Penetration enhancers, Adv Drug Del Rev, 64 (2012) 128–137.

[26] Y. Morimoto, Y. Wada, T. Seki, K. Sugibayashi, In vitro skin permeation of morphine hydrochloride during the finite application of penetration-enhancing system containing water, ethanol and l-menthol, Bio Pharm Bull, 25 (2002) 134–136.

[27] M.B. Pierre, E. Ricci, Jr., A.C. Tedesco, M.V. Bentley, Oleic acid as optimizer of the skin delivery of 5-aminolevulinic acid in photodynamic therapy, Pharm Res, 23 (2006) 360–366.

[28] F. Xie, J.K. Chai, Q. Hu, Y.H. Yu, L. Ma, L.Y. Liu, X.L. Zhang, B.L. Li, D.H. Zhang, Transdermal permeation of drugs with differing lipophilicity: Effect of penetration enhancer camphor, Int J Pharm, 507 (2016) 90–101.

[29] H. Kathuria, K. Kang, J. Cai, L. Kang, Rapid microneedle fabrication by heating and photolithography, Int J Pharm, 575 (2020) 118992.

[30] K. Ita, Transdermal drug delivery: progress and challenges, J Drug Deliv Sci Technol, 24 (2014) 245–250.

[31] D. Monti, R. Giannelli, P. Chetoni, S. Burgalassi, Comparison of the effect of ultrasound and of chemical enhancers on transdermal permeation of caffeine and morphine through hairless mouse skin in vitro, Int J Pharm, 229 (2001) 131–137.

[32] S.A. Ibrahim, S.K. Li, Efficiency of fatty acids as chemical penetration enhancers: mechanisms and structure enhancement relationship, Pharm Res, 27 (2010) 115.

[33] T. Haque, M.M.U. Talukder, enhancer: a simplistic way to modulate barrier function of the stratum corneum, Adv Pharm Bull, 8 (2018) 169–179.

[34] B. Finnin, K.A. Walters, TJ. Franz, In vitro skin permeation methodology, in: Transdermal and Topical Drug Delivery: Principles and Practice, John Wiley & Sons, 2012, pp. 85–108.

[35] L. Bartosova, J. Bajgar, Transdermal drug delivery in vitro using diffusion cells, Curr Med Chem, 19 (2012) 4671–4677.

[36] T Yano, A. Nakagawa, M. Tsuji, K. Noda, Skin permeability of various non-steroidal anti-inflammatory drugs in man, Life Sci, 39 (1986) 1043–1050.

